# Tunings for leapfrog integration of Hamiltonian Monte Carlo for estimating genetic parameters

**DOI:** 10.1101/805499

**Authors:** Aisaku Arakawa, Takeshi Hayashi, Masaaki Taniguchi, Satoshi Mikawa, Motohide Nishio

## Abstract

A Hamiltonian Monte Carlo algorithm is a Markov Chain Monte Carlo method that is considered more effective than the conventional Gibbs sampling method. Hamiltonian Monte Carlo is based on Hamiltonian dynamics, and it follows Hamilton’s equations, which are expressed as two differential equations. In the sampling process of Hamiltonian Monte Carlo, a numerical integration method called leapfrog integration is used to approximately solve Hamilton’s equations, and the integration is required to set the number of discrete time steps and the integration stepsize. These two parameters require some amount of tuning and calibration for effective sampling. In this study, we applied the Hamiltonian Monte Carlo method to animal breeding data and identified the optimal tunings of leapfrog integration for normal and inverse chi-square distributions. Then, using real pig data, we revealed the properties of the Hamiltonian Monte Carlo method with the optimal tuning by applying models including variance explained by pedigree information or genomic information. Compared with the Gibbs sampling method, the Hamiltonian Monte Carlo method had superior performance in both models. We have provided the source codes of this method written in the R language.

## Background

Computing performance has rapidly improved in recent years, and Bayesian approaches have become more popular tools for estimating genetic parameters or predicting genomic breeding values in animal and plant breeding (Meuwissen et al. 2000; Jannink et al. 2010). In particular, Bayesian inferences have been used as an alternative method of a restricted maximum likelihood (REML) method to estimate parameters if analytical models are too complicated to apply REML (Sorensen et al. 1995; Meuwissen et al. 2000; Ibáñez-Escriche et al. 2008). In Bayesian inferences, the joint posterior distribution of all parameters is constructed by multiplying a likelihood that generates the data and prior distributions, and the marginal distributions of each parameter are obtained by integrating out all other parameters of interest from the joint posterior distribution. However, the most critical limitation of an application for Bayesian approaches in quantitative genetics is that a Bayesian calculation for marginal posterior distributions often requires integration for high-dimensional distributions, and it is difficult to estimate the parameters of interest using such an analytical calculation of complex integration.

Since a series of papers of Wang et al. (1993; 1994) was published in the field of animal breeding, GS has become increasingly a popular tool for estimating genetic parameters. The GS methods have several advantages compared with the REML method, and especially, if the size of data is too large or if the models are too complex for the REML method to handle, the GS method offers, in fact, a way to estimate genetic parameters, and it can provide an effective solution by generating successive samples from conditional posterior distributions. Alternatively, in the genomic era, the GS method has been used for most Bayesian alphabet algorithms (BayesA by Meuwissen et al. (2000), BayesC by Habier et al. (2011)) for estimating single nucleotide polymorphism (SNP) effects. In most cases, the GS methods employ a single-site sampling algorithm for estimating parameters because the algorithm needs no inversion of the coefficient matrix of mixed model equations (Wang et al. 1994). Conversely, the GS method is implicitly known to require a long Markov Chain Monte Carlo (MCMC) chain to evaluate estimates of parameters of interest because the samples analyzed using the GS method are highly autocorrelated with each other, leading to a long computation time. Many researchers attempted to reduce the autocorrelations between samples using several matrix techniques under the GS scheme (García-Cortés and Sorensen 1996; Waldmann et al. 2008; Runcie and Mukherjee 2013).

Recently, the Hamiltonian Monte Carlo (HMC) algorithm has become a more popular tool in a Bayesian inference, which is based on Hamiltonian dynamics in physics (Neal 2011). The HMC algorithm was originally proposed by Duane et al. (1987) to apply the numerical simulation of lattice field. The HMC algorithm introduces an alternative variable or vector, which is called kinetic energy, to effectively transit samples within a parameter space; so, the HMC methods have a potential for giving a better sampling property than the GS methods. While the HMC algorithm could theoretically generate samples from a wide range of the parameter space with high probability, this sampling efficiency strongly depends on tunings for an approximation path integration method, a so-called leapfrog integration; the number of steps *L* and the stepsize *ϵ*. The stepsize *ϵ* governed the stability of the Hamiltonian function; for example, a larger stepsize than expected leads to a low acceptance ratio due to an increase of the integration error by the leapfrog integration. The number of steps *L* affects sampling efficiency; if *L* is not large enough, the samples generated by HMC show quite high autocorrelations between successive iterations, whereas if *L* is too large, the path approximated by leapfrog integration would retrace its previous steps of the initial state, which leads to wasted computing time (Neal 2011; Betancourt et al., 2017). Neal (2011) recommended that one practical solution for using HMC is to determine the length of trajectory, which requires selecting suitable values for *L* and *ϵ* in the leapfrog process. However, in this case, we need a lot of preliminary runs with trial values for *L* and *ϵ*, and trace plots of the preliminary runs must be checked to determine how well these runs work.

In this study, we aimed to identify the suitable values for *L* and *ϵ* for leapfrog integration to optimize the HMC method for a linear mixed model. First, we derived the HMC algorithm by applying to estimate variance components to predict breeding values, and then we searched for optimal tunings for optimizing HMC. Finally, we demonstrated the computational properties of the HMC algorithm with the optimal values of *L* and *ϵ* using real pig data.

## HMC

First, we will briefly introduce the HMC method. The HMC method is based on Hamiltonian dynamics, and the Hamiltonian (*H*) is expressed as

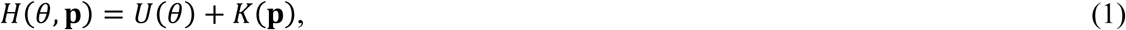

Where *U*(*θ*) and *K*(**P**) are “potential” and “kinetic” energies, respectively, in a physical system. The kinetic energy term is expressed as 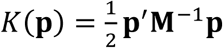, where **p** and **M** are interpreted as momentum variables and a mass matrix, respectively.

When estimating a random variable *θ* with density *p*(*θ*) using the HMC method, the independent auxiliary variable **P** is introduced. Its density is assumed to be normally distributed as follows: *p*(**p**)∼*N*(0, **M**), where **M** is interpreted as a covariance matrix in statistics. The joint density *p*(*θ*, **P**) is expressed as *p*(*θ*)*p*(**P**) because of its independence. We denoted the logarithm form of the joint density as

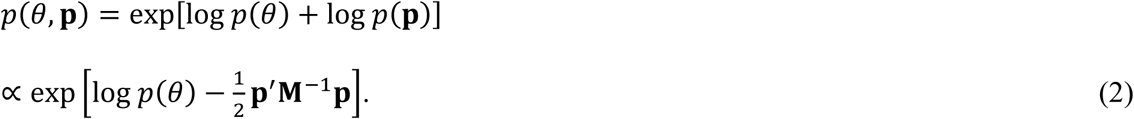

The bracketed term in equation (2) is rewritten as

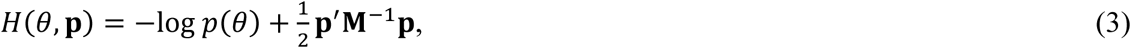

which can be interpreted as *H* with potential energy

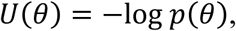

and kinetic energy

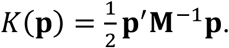

Hamilton’s equations are known as partial derivatives of *H* with respect to fictitious time *t*, and according to Neal (2011), the equations are expressed as

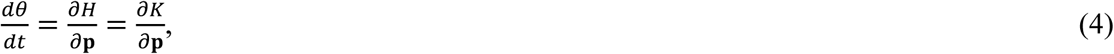

and

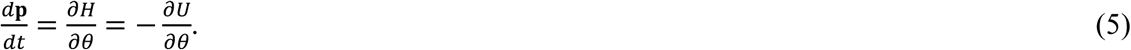

Hamiltonian dynamics has two important properties, namely, *reversibility* and *volume preservation* (Neal 2011), which rely on the use of MCMC updates. When an exact analytic solution of the differential equations (4) and (5) for Hamiltonian dynamics is available, we can use the proposed trajectory with the same volume of *H*. For practical applications, however, there is no analytic solution for Hamilton’s equations, and therefore, Hamilton’s equations must be approximated by discretizing time. The leapfrog discretization integration, also called the Stormer-Verlet method, provides a good approximation for Hamiltonian dynamics (Neal 2011). The leapfrog method depends on two arbitrarily inputted parameters, namely, *L* (the number of discrete time steps in leapfrog integration) and *ϵ* (the integration stepsize, indicating how far each leapfrog step jumps). Leapfrog integration is described as

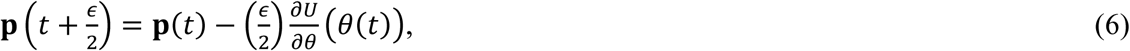

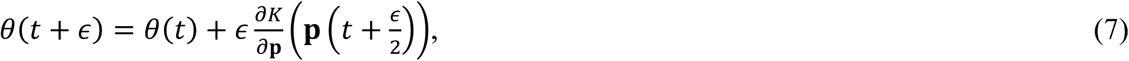

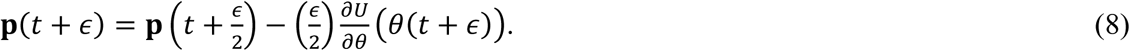

Hamiltonian dynamics is simulated using the leapfrog integration of equations (6–8) for *L* times (0 < *t* < *L*). Figure 1 shows the example for the Hamiltonian dynamics approximated by leapfrog integration. The horizontal axis is a potential variable which equals to a random variable of interest, and the vertical axis is a momentum variable sampled from a normal distribution. Each step between the consecutive *t*‘s on Figure 1 corresponds to equations 6 to 8, and *ϵ* is expressed as a difference in distance between the consecutive *t* ‘s. The preservation of volume via Hamiltonian dynamics keeps *H* invariant, but *H* is not exactly conserved with the leapfrog method because of the integration error caused by the time discretization. Therefore, a Metropolis correction step is necessary to ensure correct sampling from the marginal distribution. In the Metropolis step, the new proposal samples (*θ*\*, **P**\*) are accepted with probability

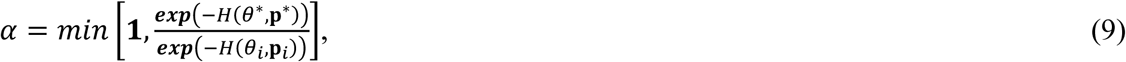

**Figure 1.**
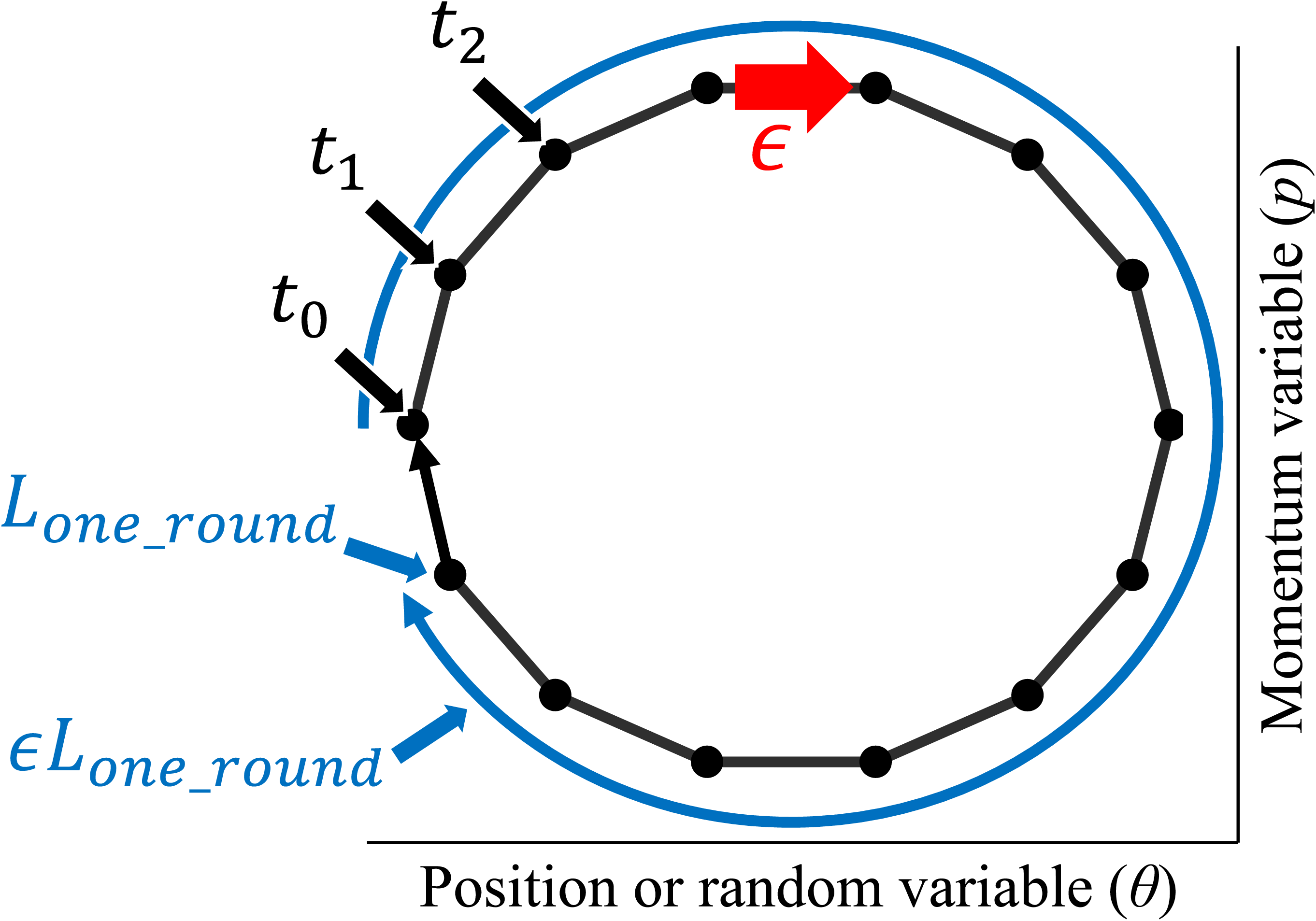
An example of a trajectory for Hamiltonian dynamics approximated by leapfrog integration. The horizontal axis shows the potential or random variable (*θ*), and the vertical axis shows the momentum variable (*p*). The stepsize is *ϵ*, and the number of step is *L* (0 1< *t* 1< *L*). The initial state is at *t*_0_, and using leapfrog integration, *θ* and *p* are moved to the next state (*t*_1_). The number of steps for one round of the trajectory is *L*_*one_round*_, and the total length of the trajectory is expressed as *ϵL*_*one_round*_.

which corresponds to the usual MH acceptance probability, or samples of (*θ*_*i*_, **P**_*i*_) keep the current (*θ*_*i*+1_, **P**_*i*+1_) = (*θ*_*i*_, **P**_*i*_). In the sampling method using Hamiltonian dynamics, *θ* and **P** are independent, and, therefore, the HMC method will give the *θ* values sampled from these marginal distributions. If the integration error in *H* remains small during the integration, then the HMC approach will achieve a high level of acceptance probability (almost 1.0).

### Linear mixed model using the HMC method

We employed a univariate linear mixed model as follows:

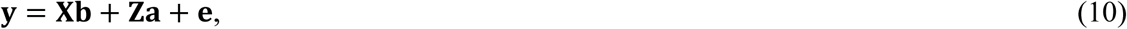

where **y** is an observation vector of order *n* × 1, **b** and **a** are location parameters with different prior distributions of orders *p* × 1 and *q* × 1, respectively, **e** is the residual error of order *n* × 1, and **X** and **Z** are designed matrices of orders *n* × *p* and *n* × *q*, respectively. The likelihood for the model and the prior distributions for **b** and **a** can be specified as 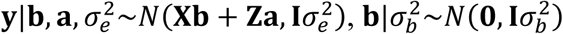 and 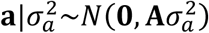, respectively, where 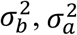 and 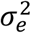 are the variances for **b, a** and **e**, respectively, **I** is an identity matrix, and **A** is a variance-covariance matrix relating to **a**. In this study, 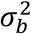 was set to be a constant value, and the prior distributions for 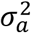 and 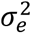 are expressed as 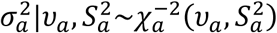 and 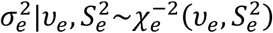, respectively, where *v*_*a*_ and *v*_*e*_ are the degrees of freedom for 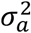 and 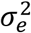, respectively, and 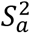 and 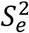 are scale parameters for 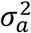 and 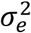, respectively. The joint distribution of parameters of the linear mixed model for the Bayesian form is expressed as 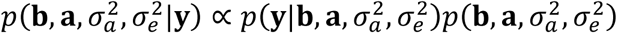, and the logarithm of the joint distribution is

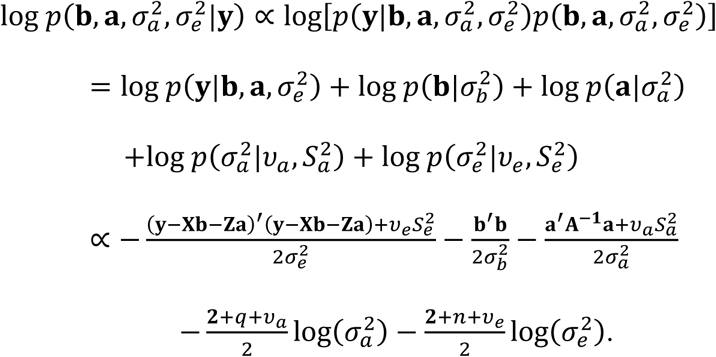

As we denote 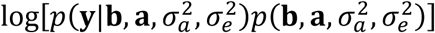 as a function *f*, we changed to

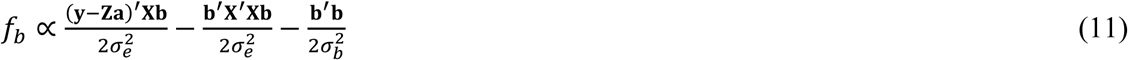

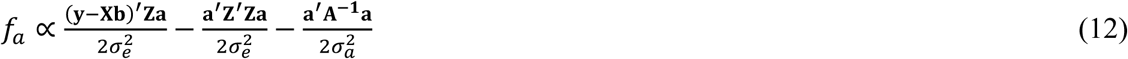

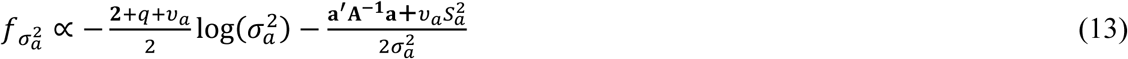

and

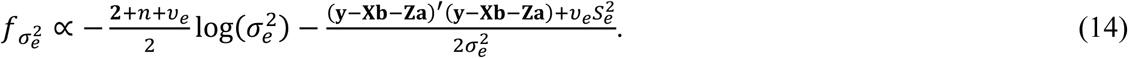

Partial derivatives for *f* with respect to each parameter (**b** or *b*_*i*_, **a** or *a*_*i*_, 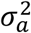, and 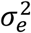, where *b*_*i*_ is the *i*th element of **b**, and *a*_*i*_ is the _*i*_th element of **a**) are expressed as follows:

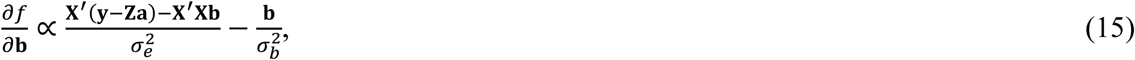

or

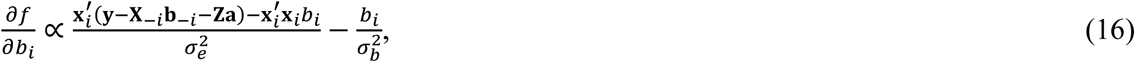

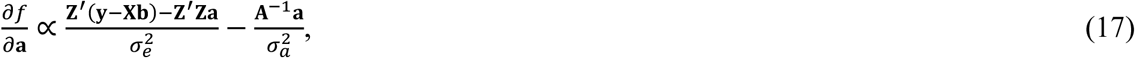

or

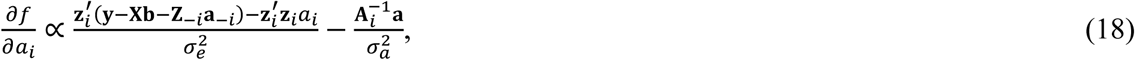

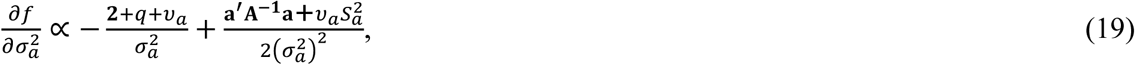

and

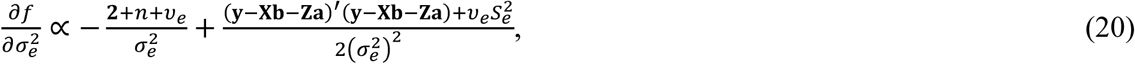

where **b**_−*i*_ is the vector of **b** without *b*_*i*_, **a**_−*i*_ is the vector of **a** without *a*_*i*_, **x**_*i*_ is the _*i*_th column vector relating to *b*_*i*_, **X**_−*i*_ is the matrix relating to **b**_−*i*_, **z**_*i*_ is the _*i*_th column vector relating to *a*_*i*_, and **Z**_−*i*_ is the matrix relating to 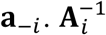 is the *i*th row vector of **A**^−1^. The HMC method could successively generate random samples from the joint posterior distribution by substituting equations (15 or 16, 17, or 18, 19, and 20) into 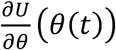 in equation (6) and 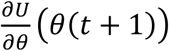 in equation (8). The pseudo-code for the HMC method with **M** = **I**, where **I** is an identity matrix, is shown in algorithm 1, and the R codes for the linear mixed model are written in Appendix III.

#### Algorithm 1. Hamiltonian Monte Carlo algorithm

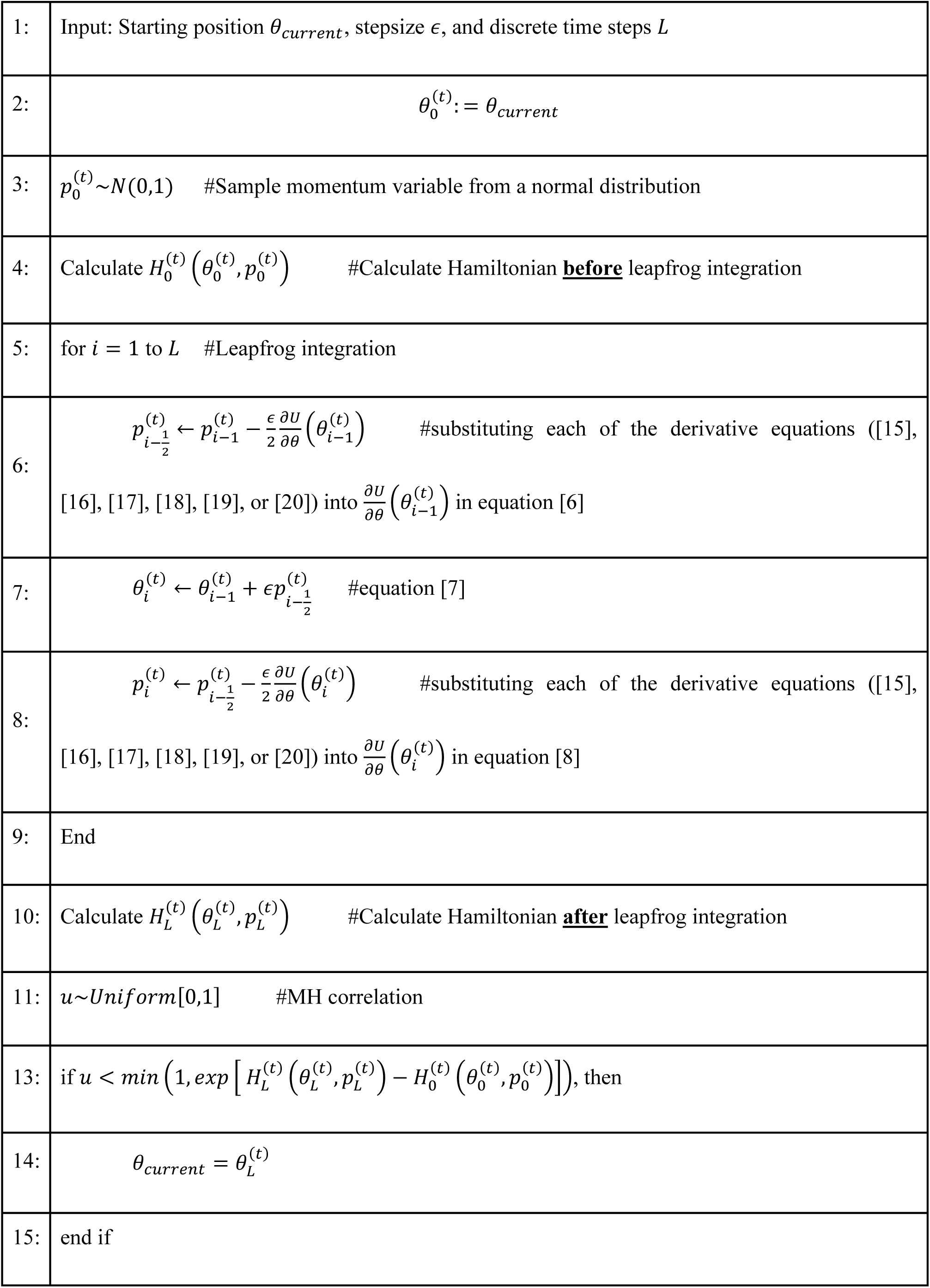

### Properties of and optimization for leapfrog integration

An HMC algorithm strongly depends on choosing two parameters of leapfrog integration; *L* and *ϵ*. In a simple situation of leapfrog integration, a trajectory of one sample shows periodically a trace on an elliptical trajectory in two dimensions (Neal 2011), as in Figure 1. We measured several steps for one round of the trajectory using leapfrog integration (*L*_*one*_*round*_ in Figure 1) to investigate the properties of the leapfrog integrations for normal and inverse chi-square distributions. A total length of the trajectory in two dimensions is expressed as *ϵL*_*one*_*round*_ (Figure 1).

Let *x* and *v* be a sample from a normal distribution *N*(*μ, σ*^2^) and a scaled inverse chi-square distribution *χ*^−1^(*n*, **u′u**), respectively, where *n* is a value of a degree of belief. Logarithm forms of these distributions are described as follows:

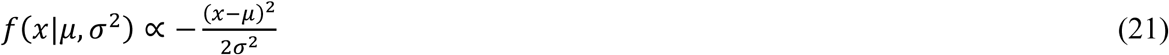

and

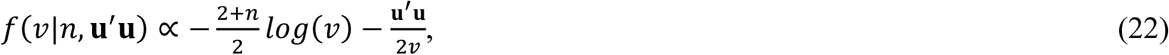

where **u′u** in equation (22) is expressed as (*n* − 1)*E*[*v*] and *E*[*v*] is an expectation value of *v*. Variances of the normal and inverse chi-square distributions are *σ*^2^ and 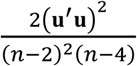, respectively. The partial derivatives for the normal and inverse chi-square distributions with respect to the parameters *x* and *v* are

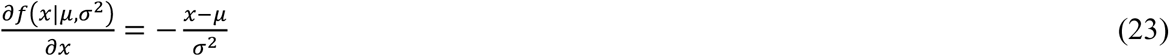

and

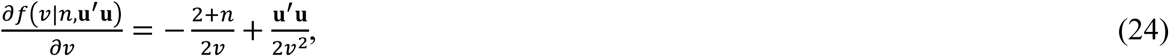

respectively. According to equations (6–8), the leapfrog integration steps might be influenced by *μ* and *σ*^2^ for the normal distribution and *n* and **u′u** for the inverse chi-square distribution. The tests were run using setting at different values of *μ* (**0** and 100) and *σ*^2^ (1, 10, and 100) for the normal distribution and different values of *n* (101, 1,001 and 10,001) and **u′u** (1,000 and 100,000) for the inverse chi-square distribution.

Rao (1945) showed the relationship between Riemann geometry and statistics, and recently, Girolami et al. (2011) incorporated a Riemann manifold in HMC, which can explain the curvature of the conditional posterior distributions by Riemann geometry. Holmes et al. (2013) and Betancourt (2017) showed geometrical interpretations for HMC. According to Riemann geometry, the Riemannian metric is defined by the Fisher information (Amari 2016). According to the similar manner of the Fisher information, second-order derivatives of the normal and the inverse chi-square distributions are

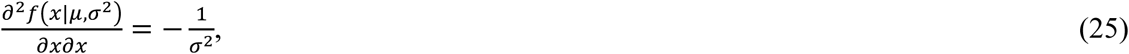

and

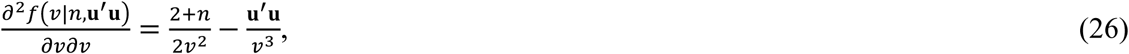

respectively. Substituting 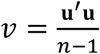 into the above equation (26), we obtained

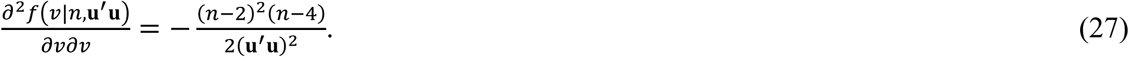

The above equations (25) and (27) correspond to negative forms of inversed variances for each distribution. In this study, we chose a square root of these variances for the two distributions as a basic indicator of *ϵ* in order to clarify the influence of the size of *ϵ* on estimation by the HMC method. *ϵ* was set to be 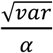, where 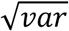 is the variance of either distribution, and *α* was set to 1, 10, and 100.

After deciding the optimal values of *ϵ*, we attempted to detect the influence of the number of *L* on the precision of estimates via the HMC method using the same simulation data by QMSim (Sargolzaei and Schenkel 2009). In the simulation, the degree of heritability was assumed to be 0.50, and phenotypic variance was set at 1.0. The base population consisted of 5 males and 50 females, and these base animals were mated at random; specifically, one male in the base population was randomly mated with 10 females to produce two males and two females of generation 1. Five males were randomly selected as sires for the next generation and mated with 10 females to produce the next generation. Five discrete generations were simulated without the base population, and the data for the base population were removed. The population size was 1,000 with equal numbers of males and females. In total, 10,000 samples were simulated, of which the first 1,000 were discarded as burn-in iterations. The post-analysis for the sampling sequences was conducted using the “*effectiveSize*” function of the R “*coda*” package (Plummer et al. 2006) to estimate the effective sample sizes (ESSs) of the sequences. We compared estimating properties for variance components and breeding values using the GS method. The starting values were set at 0.5 for the variance components and 0 for the fixed and random effects in all of the analyses. The HMC and GS programs were written in Fortran 90. In the analysis of genetic variance explained by pedigree information, we employed a sparse matrix routine by Misztal (2014).

### Application study for data

We applied the HMC method to a public pig dataset (Cleveland et al. 2012) using an infinitesimal animal model and investigated the properties of HMC sampling. In addition, we compared the HMC algorithm with GS, which is the conventional approach in Bayesian inference. According to Cleveland et al. (2012), we selected two traits that were expressed as t1 and t5 because the two traits have different genetic backgrounds; Cleveland et al. (2012) reported using the full dataset that the traits t1 and t5 are low (*h*^2^ = 0.07) and high (*h*^2^ = 0.62) heritabilities, respectively, and the phenotypic variance for t1 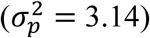 is quite lower than that for t5 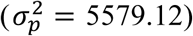.. We used phenotypic, pedigree, and genomic data. The numbers of recorded animals in t1 and t5 were 2,804 and 3,184, respectively, and pedigree information was stored for 6,473 animals. We used only SNP genotypes with a minor allele frequency of >0.05; the total number of SNPs was 45,385. We applied three single-trait models to estimate variance components; **y** = **1***μ* + **Za** + **e**, **y** = **1***μ* + **Zg** + **e**, and **y** = **1***μ* + **Zg** + **Zd** + **e**, where **a** is a vector of additive genetic effects, which is distributed 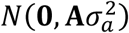; where **A** is an additive relationship matrix from pedigree information, where **g** is a vector of additive genomic effects that is distributed 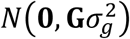, where **G** is an additive genomic relationship matrix that is calculated as in the study by VanRaden (2008), **d** is a vector of dominance deviations that is distributed 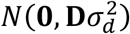, and where **D** is a covariance matrix relating to **d** that is constructed based on the work of Vitezica et al. (2013). We used only t5 for applying the model, including the dominance variance, because the previous study reported by Da et al. (2014) showed that dominance variance for t1 was quite low. Overall, 110,000 samples were simulated, the first 10,000 of which were discarded as burn-in iterations. After the samples were generated, 10,000, 50,000, and 100,000 samples after the burn-in period were used to investigate the performance of the HMC method in comparison with that of the GS method. In a post-analysis for the sampling sequences using the two methods, the “*effectiveSize*” function of the R “*coda*” package (Plummer et al. 2006) was used to estimate ESSs of the sequences.

## Results

### Leapfrog integration

Tables 1 and 2 show a summary of the number of steps for one round of the trajectory via leapfrog integration (*L*_*one*_*round*_) for the normal and the inverse chi-square distributions, respectively. For the normal distribution, when *ϵ* was expressed as the function of *σ*^2^, the scales of *μ* had no influence on the number of steps per round of the trajectory, whereas the results in Table 1 showed a linear relationship between the size of *σ*^2^ and *L*_*one*_*round*_. Consequently, the total length of the trajectory by leapfrog integration (*ϵL*_*one*_*round*_) was expressed as 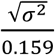. In practical application, we need the variance for conditional posterior distributions for fixed and random effects; so, Appendix I shows the deviations of *σ*^2^ for fixed and random effects.

**Table 1.** Number of steps for one round of leapfrog integration (*L*_*one*_*round*_) under the normal distribution 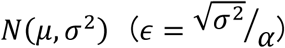

| $\alpha$ | $\mu = 0$ | | | $\mu = 100$ | | |
| --- | --- | --- | --- | --- | --- | --- |
| | $\sigma^2 = 1$ | $\sigma^2 = 10$ | $\sigma^2 = 100$ | $\sigma^2 = 1$ | $\sigma^2 = 10$ | $\sigma^2 = 100$ |
| 1 | 6 | 6 | 6 | 6 | 6 | 6 |
| 10 | 63 | 63 | 63 | 63 | 63 | 63 |
| 100 | 629 | 629 | 629 | 629 | 629 | 629 |

**Table 2.** Number of steps for one round of leapfrog integration (*L*_*oneround*_) under the inverse chi-square distribution 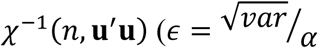, where 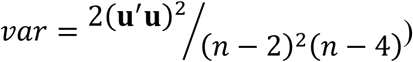

| $\alpha$ | $\mathbf{u}'\mathbf{u} = 1,000$ | | | $\mathbf{u}'\mathbf{u} = 100,000$ | | |
| --- | --- | --- | --- | --- | --- | --- |
| | $n = 101$ | $n = 1001$ | $n = 10001$ | $n = 101$ | $n = 1001$ | $n = 10001$ |
| 1 | 9 | 9 | 9 | 9 | 9 | 9 |
| 10 | 85 | 88 | 88 | 85 | 88 | 88 |
| 100 | 847 | 885 | 889 | 847 | 885 | 889 |

In the case of inverse chi-square distribution (Table 2), when the values of *ϵ* were assumed to be the function of its variance, the size of **u′u** did not affect *L*_*one*_*round*_ because the variance is a function of 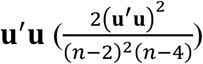, whereas the values of *n* had a little influence on *L*_*one*_*round*_, which means that if we set a higher degree for *n*, we need a longer distance of trajectory (847 in *n* = 101, and 889 in *n* = 10001 under *α* = 100). However, the effect of the values of *n* was quite small; so, we obtained the total length of the trajectory by leapfrog integration (*ϵL*_*one*_*round*_), which was expressed as 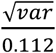, where *var* is 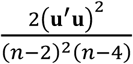.

### Inference of the discretizing time

We chose 20 on *L*_*one*_*round*_ in order to determine the optimal value for *L*, and *ϵ*s for the normal and inverse chi-square distributions were expressed as 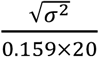 and 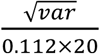. *L* was changed from 1 to 20, and the results were conducted by averaging the five different MCMC chains with a different seed. The results of the summary for variance components and breeding values for each *L* using the HMC and GS methods are shown in Table 3. The acceptance ratios were almost one in all cases, and posterior estimates using the two methods were identical to each other for all values of *L* excluding 10 and 20. The ESSs using the HMC with for *L* values of 5, 6, 8, 9, 11, 12, 13, 14, and 15 were higher than those for GS. However, in the case of *L* = _1_0, the samples of the breeding values using the HMC method had extremely high ESS values (67,267.7 ± 237,508.4). A sampling sequence of the breeding value of the animal with the largest ESS is presented in Figure 2. The graphic illustrates that the samples for the breeding value had a cyclical periodicity along the sampling sequence (Figure 2a), and that the changes of autocorrelations between lags were drastic between plus and minus (Figure 2b), suggesting that breeding values could not be sampled randomly in the case of *L* = _1_0. This nonrandom sampling for the breeding values leads to quite high estimates of genetic variance (1.39 ± 0.11).

**Table 3.**
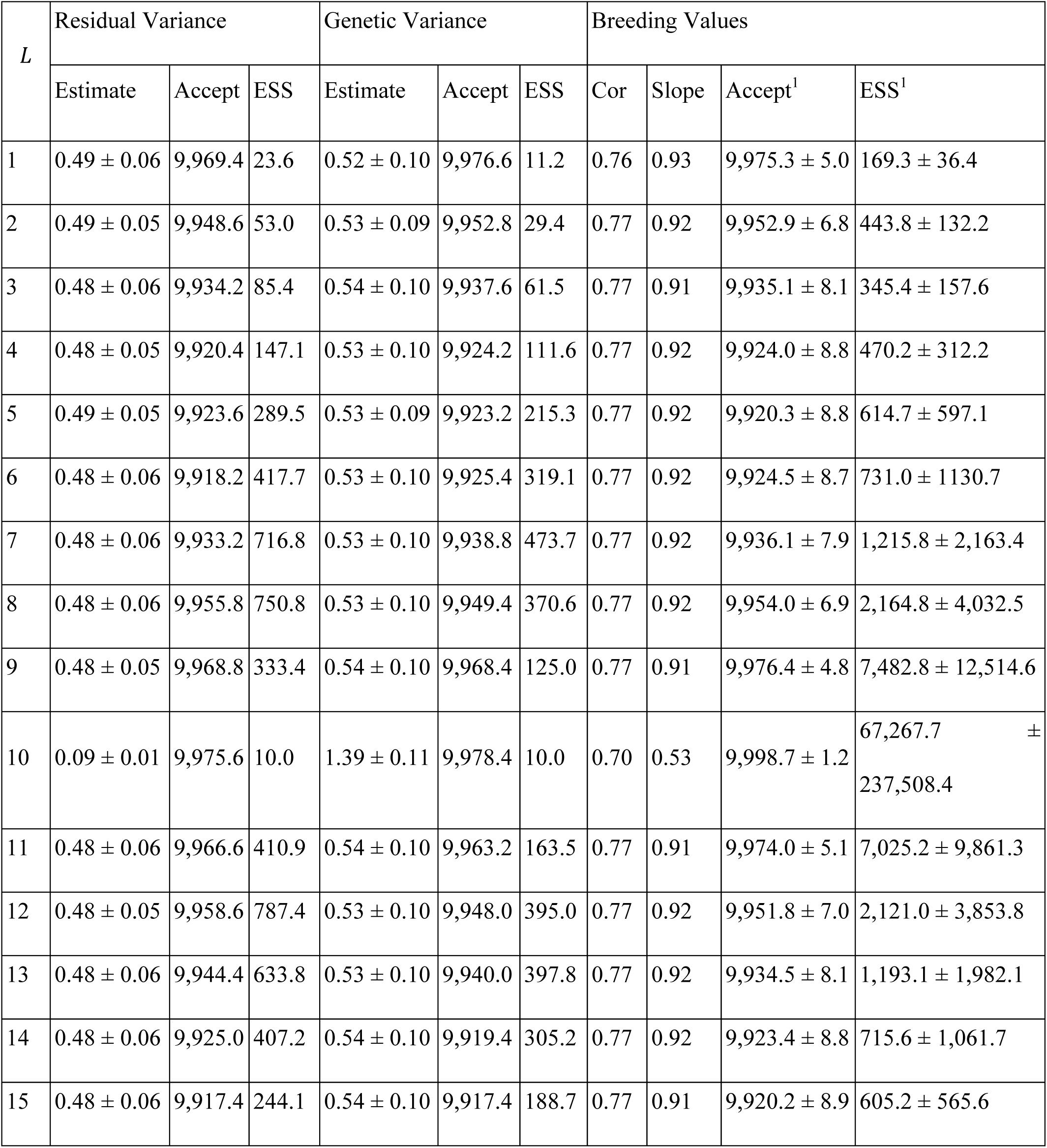
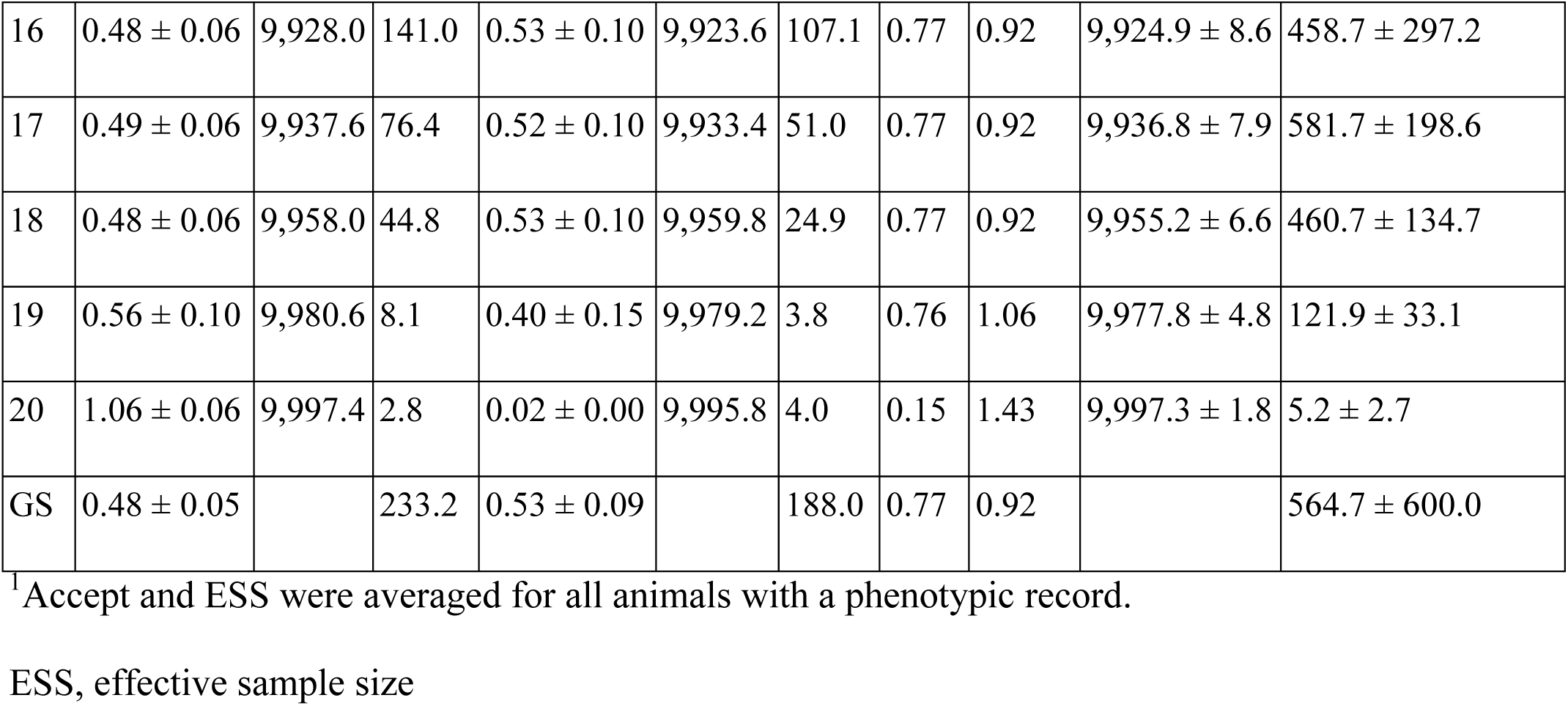
Estimates by Hamiltonian Monte Carlo (HMC) with different L values under 20 iterations per round of the trajectory for leapfrog integration and Gibbs sampling (GS) methods

**Figure 2.**
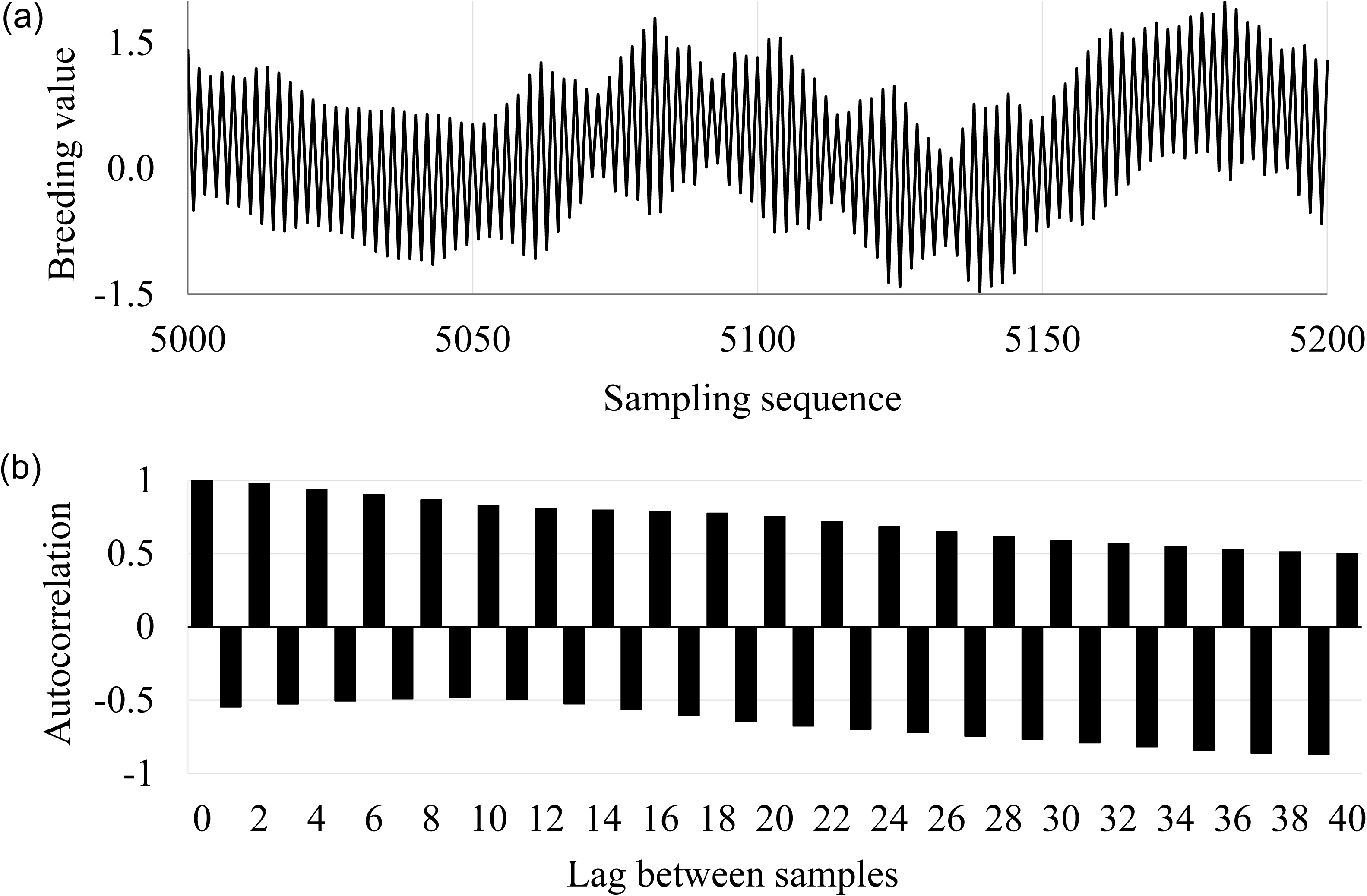
Sampling states regarding the breeding value of the phenotyped individual having the highest effective sample size. (a) The trace plot between 5,000 and 5,500. (b) Autocorrelations after the burn-in period with a sampling lag of 1–40.

### Real data

We applied the HMC method with *L* = 7 on *L*_*one*_*round*_ = 20 to the real pig data, and we also performed the analysis using 10,000, 50,000, and 100,000 samples. Summary statistics for the marginal distributions of variance components for t1 and t5 using pedigree information are shown in Tables 4 and 5, respectively, and the marginal posterior distributions for t1 and t5 are shown in Figures 3 and 4, respectively. The posterior statistics obtained using the HMC method were similar to those obtained using GS for the two traits, in line with results reported by Cleveland et al. (2012) using full pedigree information. The ESS values for the two methods generally increased linearly with an increase in the sample size. For both traits, most of the ESSs for the two variances using the HMC method were more than those using the GS method. We compared the marginal posterior distributions using the HMC method for t1 (Figure 3) and t5 (Figure 4). Excluding the genetic variances for t1, all of the variances depicted using 50,000 or 100,000 samples were similar to each other (c vs. d in Figure 3; a vs. b and c vs. d in Figure 4). In the case of 10,000 samples, the variances depicted using the HMC method were similar to those for larger sample numbers, whereas the GS method demonstrated figures that were slightly different from those for larger sample numbers. For genetic variances for t1 (Figure 3a and 3b), the marginal posterior distributions obtained using 10,000 samples exhibited polymodality and a lack of smoothness for both methods, and for the GS method, the marginal posterior distributions were also bimodal and less smooth even if 100,000 samples were generated. On the contrary, the HMC method produced a unimodal distribution that was smoother than that produced using the GS method.

**Table 4.** Summary statistics of variance components for trait 1 (t1) using the Hamiltonian Monte Carlo (HMC) and Gibbs sampling (GS) methods with 10,000, 50,000, and 100,000 sampling sequences

| Method | Sample Length | Residual Variance |  |  |  |  | Genetic Variance |  |  |  |  |
| --- | --- | --- | --- | --- | --- | --- | --- | --- | --- | --- | --- |
|  |  | Mean | Median | Mode | SD | ESS | Mean | Median | Mode | SD | ESS |
| HMC | 10,000 | 1.35 | 1.35 | 1.35 | 0.05 | 40.2 | 0.11 | 0.11 | 0.10 | 0.05 | 10.8 |
|  | 50,000 | 1.35 | 1.35 | 1.34 | 0.05 | 171.1 | 0.11 | 0.11 | 0.09 | 0.04 | 55.4 |
|  | 100,000 | 1.35 | 1.35 | 1.35 | 0.05 | 257.0 | 0.11 | 0.11 | 0.09 | 0.04 | 101.3 |
| GS | 10,000 | 1.36 | 1.36 | 1.35 | 0.03 | 69.3 | 0.11 | 0.11 | 0.11 | 0.03 | 17.2 |
|  | 50,000 | 1.37 | 1.37 | 1.37 | 0.05 | 60.9 | 0.09 | 0.09 | 0.10 | 0.04 | 23.9 |
|  | 100,000 | 1.36 | 1.36 | 1.30 | 0.05 | 142.4 | 0.10 | 0.10 | 0.07 | 0.04 | 57.6 |

ESS, effective sample size.

**Table 5.** Summary statistics of the variance components for trait 5 (t5) using the Hamiltonian Monte Carlo (HMC) and Gibbs sampling (GS) methods with 10,000, 50,000, and 100,000 sampling sequences

| Method | Sample Length | Residual Variance |  |  |  |  | Genetic Variance |  |  |  |  |
| --- | --- | --- | --- | --- | --- | --- | --- | --- | --- | --- | --- |
|  |  | Mean | Median | Mode | SD | ESS | Mean | Median | Mode | SD | ESS |
| HMC | 10,000 | 1957.6 | 1956.7 | 1962.6 | 105.9 | 366.2 | 1579.3 | 1577.9 | 1581.9 | 148.6 | 192.8 |
|  | 50,000 | 1958.5 | 1957.5 | 1963.3 | 106.7 | 1610.7 | 1576.7 | 1575.2 | 1577.1 | 148.0 | 905.9 |
|  | 100,000 | 1959.1 | 1958.0 | 1959.1 | 106.0 | 3391.1 | 1574.4 | 1572.0 | 1573.0 | 145.7 | 1871.1 |
| GS | 10,000 | 1954.8 | 1953.4 | 1961.4 | 95.5 | 204.8 | 1579.6 | 1575.7 | 1571.3 | 138.2 | 122.7 |
|  | 50,000 | 1953.5 | 1953.1 | 1953.1 | 92.1 | 976.6 | 1582.2 | 1579.0 | 1579.1 | 130.4 | 686.6 |
|  | 100,000 | 1954.0 | 1953.7 | 1954.0 | 91.5 | 1962.4 | 1581.5 | 1578.0 | 1580.1 | 128.4 | 1371.6 |

ESS, effective sample size.

**Figure 3.**
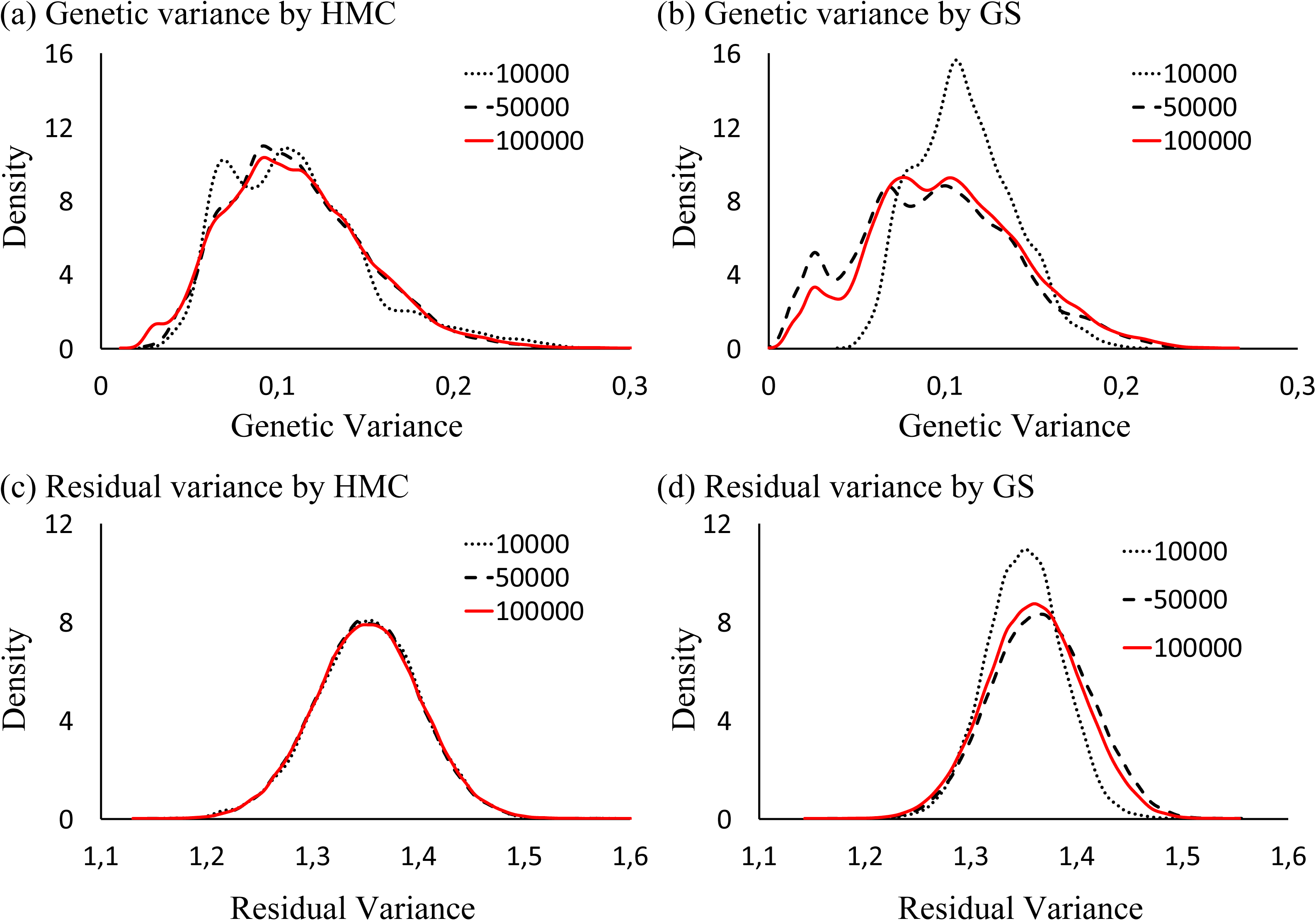
Marginal distributions of genetic 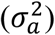 and residual 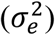 variances for t1 using the Hamiltonian Monte Carlo (HMC) and Gibbs sampling (GS) methods. (a) 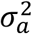 by HMC, (b) 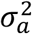 by GS, (c) 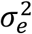 by HMC, and (d) 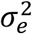 by GS.

**Figure 4.**
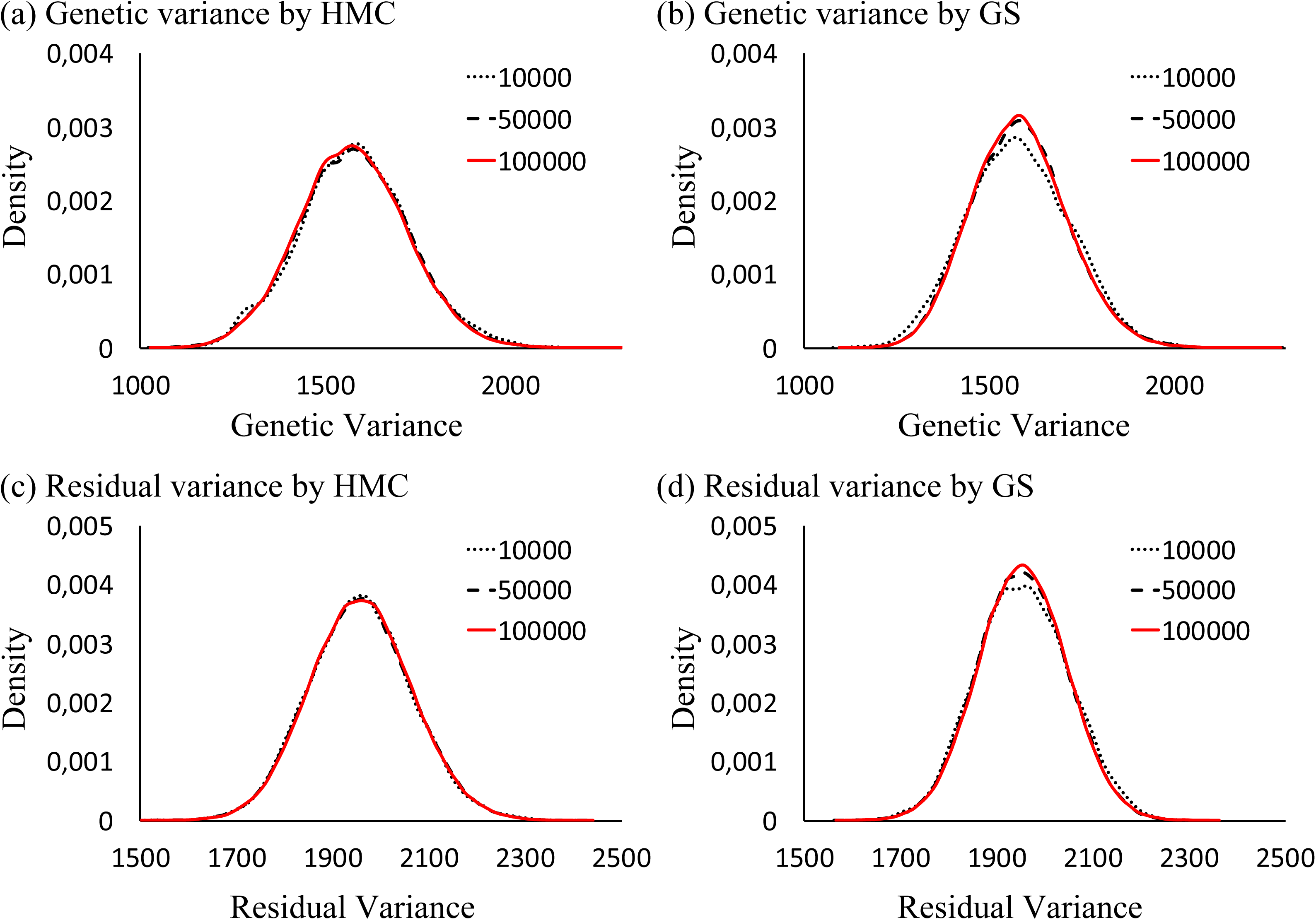
Marginal distributions of genetic 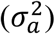 and residual 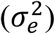 variances for t5 using the Hamiltonian Monte Carlo (HMC) and Gibbs sampling (GS) methods. (a) 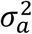 by HMC, (b) 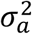 by GS, (c) 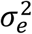 by HMC, and (d) 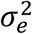 by GS.

We compared two methods applied to the same pig data but replaced the genomic information. We also performed the analysis generating 10,000, 50,000, and 100,000 samples. Summary statistics for the marginal distributions of variance components for t1 and t5 are shown in Tables 6 and 7, respectively, and the marginal posterior distributions for t1 and t5 are shown in Figures 5 and 6, respectively. The posterior statistics obtained by using the HMC method were similar to those obtained using GS for the two traits. Also, the ESS values for the two methods increased linearly with an increase in the sample size, and the ESSs for the two variances using the HMC method were much higher than those obtained by using the GS method. Compared with the marginal posterior distributions, in the trait t5, the marginal posterior distributions of all variances in the two methods were quite similar despite the sampling size (Figure 6). However, in the trait t1, the marginal distributions for genetic variances depicted using the GS method were extremely skewed even if 100,000 samples were used (b in Figure 5).

**Table 6.** Summary statistics of variance components for trait 1 (t1) using the Hamiltonian Monte Carlo (HMC) and Gibbs sampling (GS) methods with 10,000, 50,000, and 100,000 sampling sequences under genomic information.

| Method | Sample Length | Residual Variance |  |  |  |  | Genomic Variance |  |  |  |  |
| --- | --- | --- | --- | --- | --- | --- | --- | --- | --- | --- | --- |
|  |  | Mean | Median | Mode | SD | ESS | Mean | Median | Mode | SD | ESS |
| HMC | 10,000 | 1.42 | 1.43 | 1.42 | 0.04 | 134.9 | 0.03 | 0.02 | 0.02 | 0.02 | 5.1 |
|  | 50,000 | 1.42 | 1.42 | 1.42 | 0.04 | 648.1 | 0.04 | 0.03 | 0.02 | 0.02 | 36.7 |
|  | 100,000 | 1.42 | 1.42 | 1.42 | 0.04 | 1342.2 | 0.04 | 0.04 | 0.03 | 0.02 | 91.1 |
| GS | 10,000 | 1.42 | 1.42 | 1.42 | 0.03 | 177.0 | 0.04 | 0.04 | 0.03 | 0.02 | 7.7 |
|  | 50,000 | 1.44 | 1.44 | 1.44 | 0.03 | 263.2 | 0.02 | 0.01 | 0.01 | 0.02 | 11.5 |
|  | 100,000 | 1.43 | 1.43 | 1.44 | 0.03 | 350.5 | 0.02 | 0.01 | 0.01 | 0.02 | 24.8 |

ESS, effective sample size.

**Table 7.** Summary statistics of the variance components for trait 5 (t5) using the Hamiltonian Monte Carlo (HMC) and Gibbs sampling (GS) methods with 10,000, 50,000, and 100,000 sampling sequences under genomic information.

| Method | Sample Length | Residual Variance |  |  |  |  | Genomic Variance |  |  |  |  |
| --- | --- | --- | --- | --- | --- | --- | --- | --- | --- | --- | --- |
|  |  | Mean | Median | Mode | SD | ESS | Mean | Median | Mode | SD | ESS |
| HMC | 10,000 | 2161.2 | 2160.0 | 2155.9 | 76.1 | 1165.6 | 1322.7 | 1315.6 | 1301.5 | 126.9 | 267.2 |
|  | 50,000 | 2161.4 | 2160.4 | 2153.3 | 75.6 | 6366.6 | 1318.9 | 1314.5 | 1299.5 | 122.1 | 1600.4 |
|  | 100,000 | 2161.7 | 2160.7 | 2154.9 | 75.4 | 12142.9 | 1318.1 | 1314.0 | 1293.7 | 123.3 | 3086.1 |
| GS | 10,000 | 2160.7 | 2160.1 | 2155.5 | 64.4 | 435.7 | 1316.5 | 1309.4 | 1300.6 | 119.3 | 183.0 |
|  | 50,000 | 2159.4 | 2158.8 | 2153.6 | 63.5 | 2388.0 | 1321.0 | 1315.3 | 1294.8 | 113.6 | 970.3 |
|  | 100,000 | 2161.3 | 2160.9 | 2156.9 | 63.4 | 4519.7 | 1312.1 | 1312.1 | 1293.6 | 113.3 | 1973.0 |

ESS, effective sample size.

**Figure 5.**
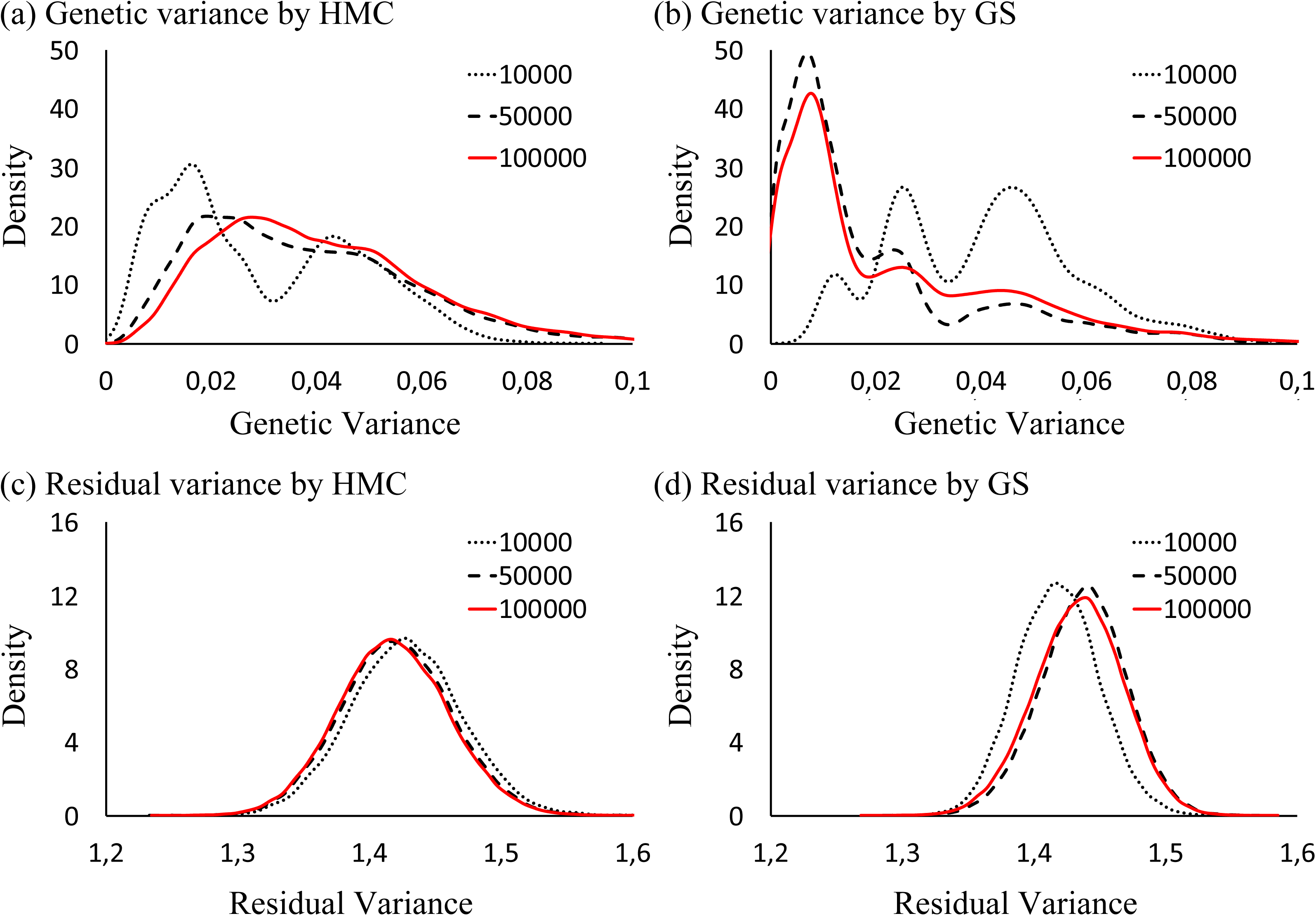
Marginal distributions of genomic 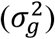 and residual 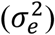 variances for t1 using the Hamiltonian Monte Carlo (HMC) and Gibbs sampling (GS) methods. (a) 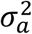 by HMC, (b) 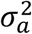 by GS, (c) 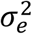 by HMC, and (d) 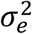 by GS.

**Figure 6.**
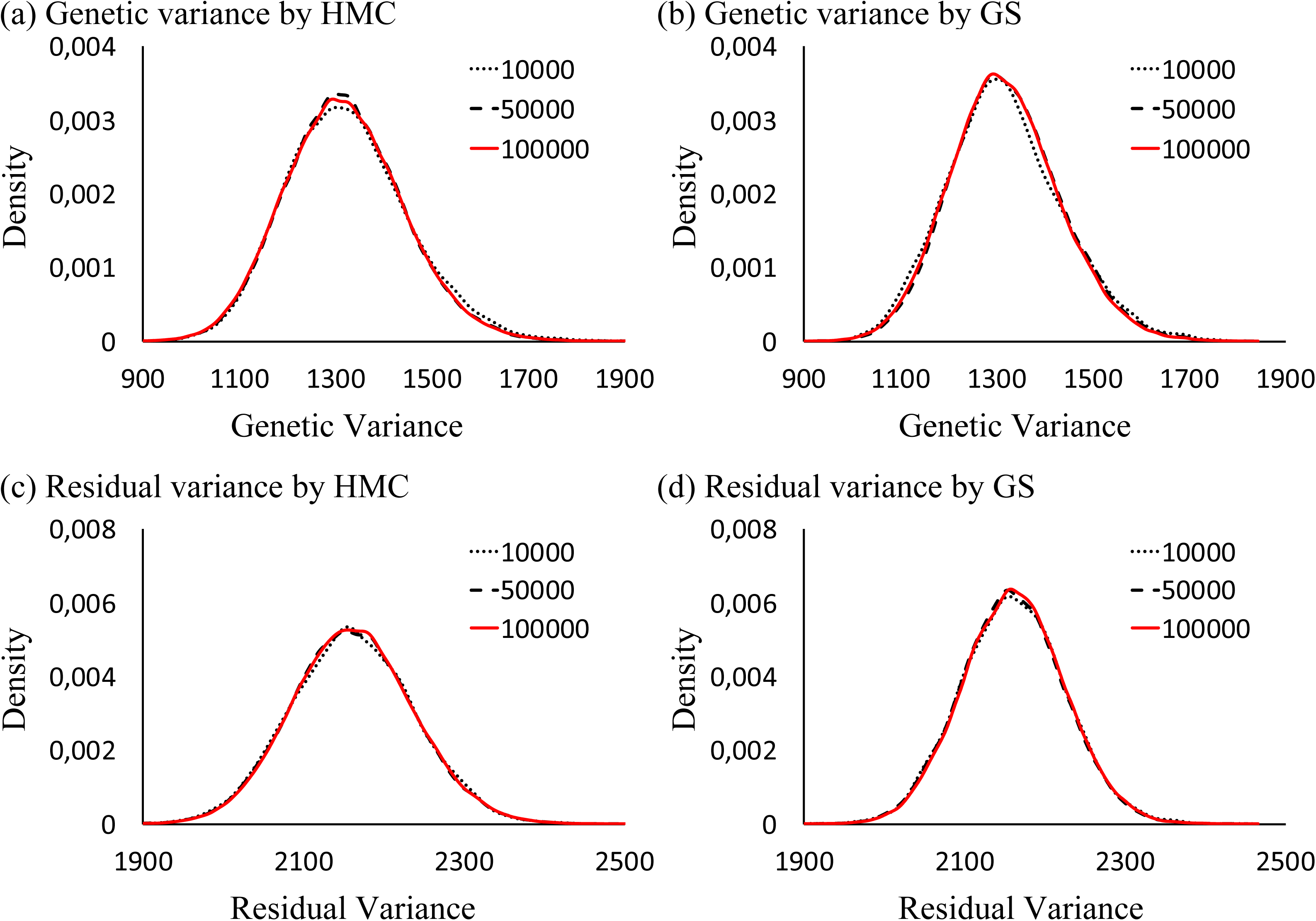
Marginal distributions of genomic 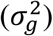 and residual 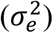 variances for t5 using the Hamiltonian Monte Carlo (HMC) and Gibbs sampling (GS) methods. (a) 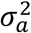 by HMC, (b) 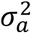 by GS, (c) 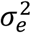 by HMC, and (d) 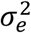 by GS.

In more complex situations, such as in the models including non-additive genetic effects like dominance deviations, the summary statistics for the marginal distributions of variance components for t5 are shown in Table 8, and the marginal posterior distributions are shown in Figure 7. In the dominance variance, using the HMC method produced similar estimates despite the sample size; however, the estimates of the GS method appeared unstable while generating 100,000 samples and were lower than those obtained using the HMC method. The marginal distributions of the dominance variance using the GS method were slightly skewed compared with those obtained using the HMC method.

**Table 8.** Summary statistics of variance components for trait 1 (t5) using the Hamiltonian Monte Carlo (HMC) and Gibbs sampling methods with 10,000, 50,000, and 100,000 sampling sequences

| Method | Sample Length | Residual Variance |  |  |  |  | Additive Genomic Variance |  |  |  |  | Dominance Genomic variance |  |  |  |  |
| --- | --- | --- | --- | --- | --- | --- | --- | --- | --- | --- | --- | --- | --- | --- | --- | --- |
|  |  | Mean | Median | Mode | SD | ESS | Mean | Median | Mode | SD | ESS | Mean | Median | Mode | SD | ESS |
| HMC | 10,000 | 1981.3 | 1981.4 | 1975.7 | 87.3 | 154.3 | 1313.5 | 1309.3 | 1293.2 | 125.7 | 281.5 | 186.4 | 178.9 | 166.6 | 57.7 | 28.0 |
|  | 50,000 | 1982.6 | 1981.0 | 1975.7 | 91.4 | 602.6 | 1308.3 | 1304.2 | 1291.5 | 122.7 | 1469.1 | 186.1 | 182.8 | 167.2 | 65.0 | 93.1 |
|  | 100,000 | 1986.6 | 1985.0 | 1980.9 | 92.3 | 1033.2 | 1306.4 | 1302.2 | 1289.0 | 123.7 | 3077.6 | 182.8 | 180.6 | 172.7 | 66.2 | 201.3 |
| Gibbs sampling | 10,000 | 2007.4 | 2008.2 | 2008.4 | 80.9 | 78.9 | 1302.3 | 1299.5 | 1298.3 | 111.4 | 193.2 | 156.7 | 155.2 | 160.6 | 63.4 | 13.1 |
|  | 50,000 | 2022.6 | 2021.6 | 2021.8 | 82.5 | 324.9 | 1296.6 | 1294.6 | 1289.6 | 111.0 | 972.6 | 144.0 | 143.7 | 161.1 | 63.1 | 59.0 |
|  | 100,000 | 2010.0 | 2008.8 | 2002.5 | 83.2 | 688.5 | 1300.9 | 1297.8 | 1293.6 | 114.1 | 1827.3 | 156.3 | 157.4 | 166.5 | 63.1 | 129.2 |

ESS, effective sample size.

**Figure 7.**
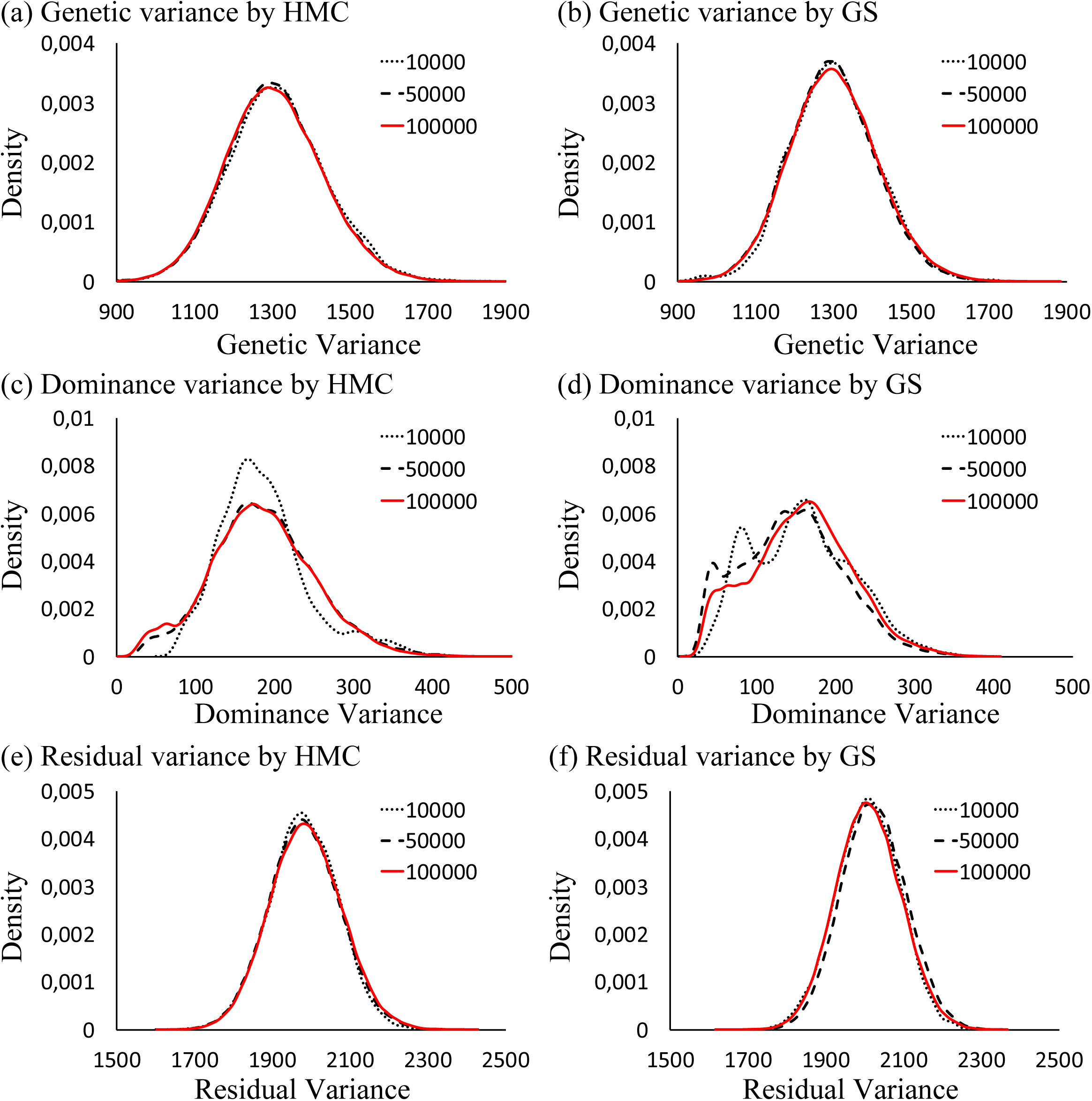
Marginal distributions of additive genomic 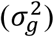, dominance genomic 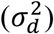, and residual 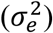 variances for t5 using the Hamiltonian Monte Carlo (HMC) and Gibbs sampling (GS) methods. (a) 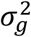 by HMC, (b) 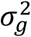 by GS, (c) 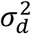 by HMC, (d) 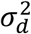 by GS, (e) 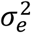 by HMC, and (f) 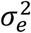 by GS.

We compared the computation times of both methods, using MacBook pro on an Intel Core i7 Processor (2.7 GHz) with 16 GB of RAM. We generated five chains with different seeds for the trait t1 using the model, including the genomic additive variance. In HMC, each simulation took 5,957.8 ± 12.6 sec, while the corresponding average time for GS was 5,924.1 ± 4.1 sec. In the HMC iteration, the leapfrog integrations of fixed and random effects are decomposed into the right and left hands of Henderson’s mixed model equations (Appendix II), which include several matrix-vector multiplications. The same multiplications are needed in the GS iteration. These matrix-vector multiplication calculations are heavier than the leapfrog integration in HMC or a random number generator in GS. Consequently, the total computational time is quite similar to each other.

## Discussion

Bayesian statistics have provided large amounts of information, and the GS method is the conventional Bayesian approach in animal breeding, being a feasible procedure for constructing the posterior distribution of interest. In this study, we proposed another MCMC method, called HMC, which is based on Hamilton dynamics, for estimating genetic parameters in the animal breeding fields. The HMC method also requires consideration of sampling convergence, length of the burn-in period, the number of samples, and ESS similar to other MCMC methods. Recently, many complex models, such as random regression models (Jamrozik and Schaeffer 1997) as well as a single-step genomic best linear unbiased prediction (Aguilar et al. 2010), have been proposed for animal breeding analyses. The likelihood functions of these analyses are likely too complex to be performed using REML. Although the GS method can provide marginal posterior distributions, in the case of analyzing complex models for which REML is inapplicable, GS exhibits extremely slow convergence and generates highly autocorrelated samples. Contrarily, the HMC method uses gradient information about a logarithm of a posterior distribution to investigate the distribution space, which may lead to better mixing properties than the MH and GS methods.

The HMC method is an efficient sampling method, but the sampling performance of the HMC method strongly depended on the leapfrog integration parameters *L* and *ϵ* (Neal et al. 2011). The leapfrog integration process given a relatively large *ϵ* could not approximate the path for the trajectory adequately during the discretization time, and if using a small *ϵ*, more time would be needed to approximate the distance of a trajectory via leapfrog integration. If we set the same *L* in the HMC method, we could not obtain samples from marginal distribution under the large *ϵ*, while the samples are similar to those of the previous iteration under the small *ϵ*. In our study, we tried to reveal the properties of leapfrog integration for the normal and inverse chi-square distributions. When one parameter is considered, a trajectory of an approximate path using leapfrog integration according to equations (6–8) shows an ellipse on two dimensions. In order to optimize the performance of the HMC method, we decided on the length of the trajectory of the normal and the inverse chi-square distributions as the value of *ϵL*_*one_round*_. When *L*_*one_round*_ was constant at 20 as a maximum discretization time, an *L* value of 7 provided a good performance for the HMC method. However, our settings for the HMC method were only used in the model, including random effects with no correlations. Therefore, we would need a modification to handle the correlation parameters, such as genetic correlations.

The parameter space of a statistical model can be expressed as a Riemann manifold, which can define the structure of the posterior distribution geometrically (Rao 1945). Girolami and Calderhead (2011) showed a more excellent way of incorporating the MHC algorithm into Riemann geometry in order to address many of the shortcomings of HMC. This algorithm is called Riemannian Manifold HMC (RMHMC), which can explain the curvature of the conditional posterior distributions by Riemann geometry. In this theory, an information matrix **G**(*θ*) is used instead of a fixed mass matrix **M** of the kinetic energy term *K*(**P**), and the kinetic energy term is modified as

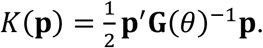

Girolami and Calderhead (2011) used the expected Fisher information matrix as **G**(*θ*), which is defined as a positive semidefinite, whereas Paquet and Fraccaro (2016) used the observed Fisher information matrix. Although our results partially related to the Riemannian manifold, compared with these studies, we assigned the square root of variances of the conditional distributions to *ϵ* rather than **M** of the kinetic energy term. In addition, the variances of the conditional distributions do not correspond completely to the Fisher information. Our approach projected Hamiltonian function onto a Euclidean manifold and did not fully consider the local structure of the target distribution. Therefore, our approach would not be able to fully guarantee sampling from the marginal distributions within a parameter space when true values are on the edge of the parameter space. We applied the HMC with our tunings to extreme simulated data (10 individuals, *h*^2^ = 0, and 10,000 SNP markers with equilibrium). We generated 10 different datasets with different seeds. We analyzed using the model including an additive and dominance genomic variance. As a result, the HMC method with our tunings did not fail outside of the parameter spaces for the variance components and breeding values, suggesting that our turnings could estimate parameters on the edge of the parameter space without failure (data not shown).

Our approach has two advantages compared with the RMHMC algorithm. First, it was easy to apply the HMC algorithm to a single trait linear mixed model because we only use 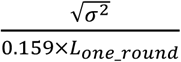 and 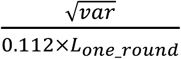 for the normal and the inverse chi-square distributions, respectively, as *ϵ*. Second, in the RMHMC algorithm, the Fisher information and a first-order derivative of the Fisher information are needed in the leapfrog process. Therefore, potential energy is no longer independent from kinetic energy in RMHMC; hence, fixed point iterations must be employed in the process of leapfrog integration of RMHMC, which suggests that RMHMC needs more nested iterations within leapfrog integration.

In comparison with the computing times of the GS methods, theoretically, the HMC method requires a longer computing time for the discrete time steps (*L*) than the GS method, but the HMC method showed similar computing time with the GS method in the context of genomic analysis. As previously mentioned, in the context of the mixed model, both HMC and GS require the same times of matrix-vector multiplications in the sampling of fixed and random effects in each iteration, which involves heavy computation within MCMC iterations.

The HMC method gave a higher ESS compared with the GS method, and the samples from the HMC methods could be generated from a wider range of its sampling space. Therefore, it would be possible for the HMC methods to shorten the total sample size, which leads to markedly decreasing the total computing time in HMC. Furthermore, in this study, the HMC method showed better sampling properties in the case of low heritability than those of the GS method because the HMC method gave a relatively smooth marginal distribution even in low heritability (the additive genetic variances in Figures 3 and 5 and the dominance genetic variances in Figure 7).

Many HMC algorithms have been developed to be free of problems concerning leapfrog integration and to shorten the burn-in period or accelerate its mixing properties. The most popular algorithm is No-U-Turn-Sampler (NUTS) (Hoffman and Gelman 2014), and STAN software (Carpenter et al. 2017), which has rapidly gained popularity in many Bayesian analysis fields, is equipped with this algorithm. The NUTS algorithm is extremely effective for the sampling process because it automates tuning in leapfrog integration, as neither the step size nor the number of steps needs to be specified by the user. However, the algorithm has a severe disadvantage regarding computing time because NUTS needs to construct a deep binary tree in each step to specify an optimal *L* value. Additionally, STAN is a stand-alone program, and thus, it is difficult to modify it to analyze large amounts of animal breeding data as those generated for estimating genomic evaluation. Compared with our optimized HMC method, we need to set the leapfrog tuning beforehand, and the numbers of steps and stepsizes on our algorithm are not determined successively for each transition in each iteration.

In this study, we developed the HMC algorithm for a simple mixed model and optimized the algorithm to enable effective sampling from marginal posterior distributions. HMC could be generalized to more complex situations, such as a multiple-traits model (Van Tassel and Van Vleck 1996) or a threshold model (Sorensen et al. 1995), but we need to identify another optimization parameter of leapfrog integration for covariance components or thresholds; contrarily, a more flexible algorithm, such as RMHMC, must be applied to these generalized models.

## Conclusion

In this study, we examined the HMC algorithm in the context of a linear mixed mode on quantitative genetics. This method strongly depends on two parameters, *ϵ* and *L*, for leapfrog integration. We applied one of the tunings for the integration process. The HMC method with optimized tuning provided superior sampling performances compared to the GS method. In addition, the HMC method appeared to generate samples from a wider range of parameter spaces than the GS method. The complete R and Fortran scripts are available from Aisaku Arakawa on reasonable request.

## Acknowledgments

The authors thank Dr. Andres Legarra at INRA Toulouse for his constructive comments on an earlier manuscript version. A.A. conducted a portion of this work while visiting INRA Toulouse.

## Appendixes

### Appendix I

According to Wang et al. [10], the factorization form of th fixed effect 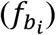 and *i*th breeding values 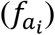 can be expressed as

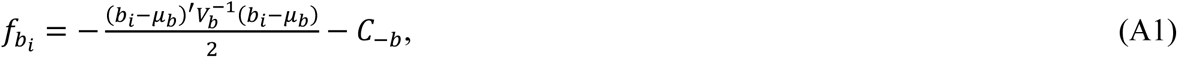

and

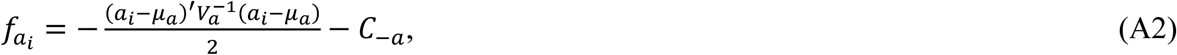

where *b*_*i*_ and *a*_*i*_ are the *i*th fixed effect and *i*th breeding value, respectively, *C*_−*b*_ and *C*_−*a*_ are components not including elements related to *b*_*i*_ and *a*_*i*_, respectively, *μ*_*b*_ and *μ*_*a*_ are described as

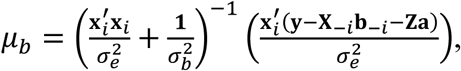

and

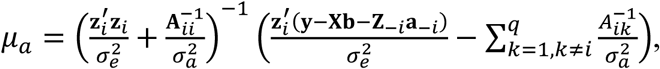

respectively, and *V*_*b*_ and *V*_*a*_ are described as

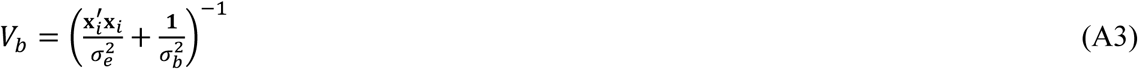

and

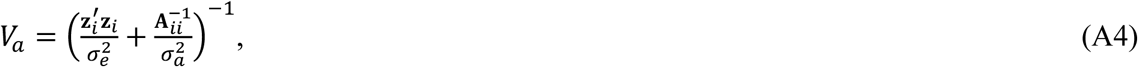

respectively. The two factorization forms (A1) and (A2) can be regarded as normal distributions of *b*_*i*_|*μ*_*b*_, *V*_*b*_∼*N*(*μ*_*b*_, *V*_*b*_) and *a*_*i*_|*μ*_*a*_, *V*_*a*_∼*N*(*μ*_*a*_, *V*_*a*_).

### Appendix II

We can change equations (15–18) of leapfrog integration, and the equations are expressed as follows:

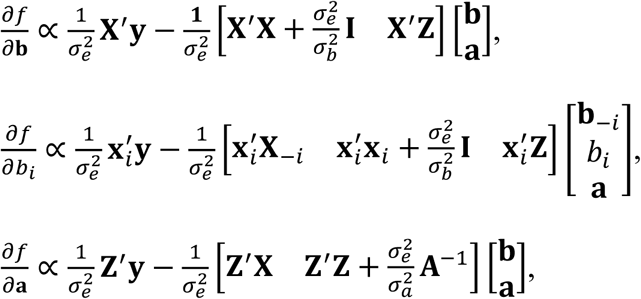

and

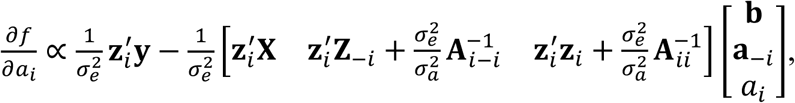

where **b**_−*i*_ is the vector of **b** without *b*_*i*_, **a**_−*i*_ is the vector of **a** without *a*_*i*_, **x**_*i*_ is the *i*th column vector relating to *b*_*i*_, **X**_−*i*_ is the matrix relating to **b**_−*i*_, **z**_*i*_ is the *i*th column vector relating to *a*_*i*_, and **Z**_−*i*_ is the matrix relating to 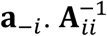 is the scalar value of the *i*th row and *i*th column of **A**^−1^, and 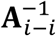 is the *i*th row vector of **A**^−1^ without 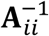. In each equation, the first terms of the right hands are the right hands of the Henderson’s mixed model equations, and the second terms are the left hands of the mixed model equations.

### Appendix III

Appendix III demonstrates R codes for the HMC method. Table A1 shows a brief description of the variables we used in our R code.

**Table A1.** Variables used in our R codes.

|  |  |  |
| --- | --- | --- |
| Basic parameters | num.p (**constant**) | a total size of fixed effect |
|  | num.ped (**constant**) | the number of pedigrees |
|  | n (**constant**) | the number of phenotypes |
| Model description | y (**constant**) | phenotypic vector (n) |
|  | b | fixed effect work vector (num.p) |
|  | u | random effect (breeding values) work vector (size num.ped) |
|  | X (**constant**) | designed matrix for b (n, num.p) |
|  | Z (**constant**) | designed matrix for u (n, num.ped) |
|  | A.inv (**constant**) | a matrix of the inverse of an additive relationship matrix (num.ped, num.ped) |
| Variances | tau (**constant**) | prior variance for b, we set tau = 10,000 |
|  | var.u | genetic variance work variable |
|  | var.e | residual variance work variable |
| Computing efficiency vector | xx (**constant**) | diagonal elements for $\mathbf{X}'\mathbf{X}$ (num.p) |
| | zz (**constant**) | diagonal elements for $\mathbf{Z}'\mathbf{Z}$ (num.ped) |
| Leapfrog integration | epsilon | $\epsilon$ |
| | L (**constant to 7**) | $L$ |
| | max.L (**constant to 20**) | $L_{one\_round}$; maximum iterations per one round |

### 1) R code for the HMC method of fixed effects

L <- 7; max.L <- 20

e <- y-X%*%b-Z%*%u

for(i in 1:num.p){

epsilon <- sqrt(1/(xx[ii]/var.e+1/tau))/(0.1589825*max.L)

b.tmp <- b[i]

xe <- crossprod(e+X[,i]*b.tmp, X[,i])

xx <- crossprod(X[,i])

p <- rnorm(1)

K0 <- t(p)%*%p/2

U0 <- -((2*xe*b.tmp-xx[i]*b.tmp^2)/(2*var.e)-b.tmp^2/(2*tau))

H0 <- (U0+K0)

for(t in 1:L){

p <- p-0.5*epsilon*(-((xe-xx[jj]*b.tmp)/var.e-b.tmp/tau))

b.tmp <- b.tmp+epsilon*p

p <- p-0.5*epsilon*(-((xe-xx[jj]*b.tmp)/var.e-b.tmp/tau))

}

K1 <- t(p)%*%p/2

U1 <- -((2*xe*b.tmp-xx[jj]*b.tmp^2)/(2*var.e)-b.tmp^2/(2*tau))

H1 <- (U1+K1)

if(runif(1) > exp(H0-H1)) b.tmp <- b[i]

e <- e+X[,i]*c(b[i]-b.tmp)

b[i] <- b.new

}

### 2) R code for the HMC method of breeding values

e <- y-X%*%b-Z%*%u

for(i in 1:num.ped){

epsilon <- sqrt(1/(zz[jj]/var.e+A.inv[jj,jj]/var.u))/(0.1589825*max.L)

u.tmp <- u[i]

ze <- crossprod(e+Z[,i]*u.tmp, Z[,i])

zz <- crossprod(Z[,i])

uG <- crossprod(A.inv[,i],u)-A.inv[i,i]*u.tmp

p <- rnorm(1)

K0 <- t(p)%*%p/2

U0 <- -((2*ze*u.tmp-zz[i]*u.tmp^2)/(2*var.e)-(2*uG*u.tmp+A.inv[i,i]*u.tmp^2)/(2*var.u))

H0 <- (U0+K0)

for(t in 1:L){

p <- p-0.5*epsilon*(-((ze-zz[i]*u.tmp)/var.e-(uG+A.inv[i,i]*u.tmp)/var.u))

u.tmp <- u.tmp+epsilon*p

p <- p-0.5*epsilon*(-((ze-zz[i]*u.tmp)/var.e-(uG+A.inv[i,i]*u.tmp)/var.u))

}

K1 <- t(p)%*%p/2

U1 <- -((2*ze*u.tmp-zz[i]*u.tmp^2)/(2*var.e)-(2*uG*u.tmp+A.inv[i,i]*u.tmp^2)/(2*var.u))

H1 <- (U1+K1)

if(runif(1) > exp(H0-H1)) u.tmp <- u[i]

e <- e+Z[,i]*c(u[i]-u.tmp)

u[i] <- u.new

}

### 3) R code for the HMC method of genetic variance

var.u.tmp <- var.u

uAu <- t(u)%*%a.inv%*%u

epsilon <- sqrt(uAu**2/((num.ped-1)^2*(num.ped-2)))/(0.112485939* max.L)

p <- rnorm(1)

K0 <- t(p)%*%p/2

U0 <- -(-((num.ped+v.u)/2+1)*log(var.u.tmp)-(uAu+lambda.u)/(2*var.u.tmp))

H0 <- (U0+K0)

for(t in 1:L){

p <- p-0.5*epsilon*(-(-((num.ped+v.u)/2+1)/var.u.new+

0.5*(uAu+lambda.u)/(var.u.tmp^2)))

var.u.tmp <- var.u.tmp+epsilon*p

p <- p-0.5*epsilon*(-(-((num.ped+v.u)/2+1)/var.u.tmp+

0.5*(uAu+lambda.u)/(var.u.tmp^2)))

}

K1 <- t(p)%*%p/2

U1 <- -(-((num.ped+v.u)/2+1)*log(var.u.tmp)-(uAu+lambda.u)/(2*var.u.tmp))

H1 <- (U0+K0)

if(runif(1) < exp(-H1+H0)) var.u <- var.u.tmp

### 4) R code for the HMC method of residual variance

var.e.tmp <- var.e

e <- y-X%*%b-Z%*%u

ee <- t(e)%*%e

epsilon <- sqrt(ee**2/((n-1)^2*(n-2)))/(0.112485939* max.L)

p <- rnorm(1)

K0 <- t(p)%*%p/2

U0 <- -(-((n+v.e)/2+1)*log(var.e.tmp)-(ee+lambda.e)/(2*var.e.tmp))

H0 <- (U0+K0)

for(t in 1:L){

p <- p-0.5*epsilon*(-(-((n+v.e)/2+1)/var.e.tmp+

0.5*(ee+lambda.e)/(var.e.tmp^2)))

var.e.tmp <- var.e.tmp+epsilon*p

p <- p-0.5*epsilon*(-(-((n+v.e)/2+1)/var.e.tmp+

0.5*(ee+lambda.e)/(var.e.tmp^2)))

}

K1 <- t(p)%*%p/2

U1 <- -(-((n+v.e)/2+1)*log(var.e.new)-(ee+lambda.e)/(2*var.e.new))

H1 <- (U1+K1)

if(runif(1) < exp(H0-H1)) var.e <- var.e.new

